# Premature translation of the zygotic genome activator Zelda is not sufficient to precociously activate gene expression

**DOI:** 10.1101/2022.03.22.485419

**Authors:** Elizabeth D. Larson, Hideyuki Komori, Zoe A. Fitzpatrick, Samuel D. Krabbenhoft, Cheng-Yu Lee, Melissa Harrison

## Abstract

Following fertilization, the unified germ cells rapidly transition to a totipotent embryo. Maternally deposited mRNAs encode the proteins necessary for this reprogramming as the zygotic genome remains transcriptionally quiescent during the initial stages of development. The transcription factors required to activate the zygotic genome are among these maternally deposited mRNAs and are robustly translated following fertilization. In *Drosophila*, the mRNA encoding Zelda, the major activator of the zygotic genome, is not translated until one hour after fertilization. Here we demonstrate that *zelda* translation is repressed following fertilization by the TRIM-NHL protein Brain tumor (BRAT). BRAT also regulates Zelda levels in the larval neuroblast lineage. In the embryo, BRAT-mediated translational repression is regulated by the Pan Gu (PNG) kinase, which is triggered by egg activation. The PNG kinase phosphorylates translational regulators, suggesting that PNG kinase activity alleviates translational repression of *zelda* by BRAT and coupling translation of *zelda* with that of other regulators of early embryonic development. Using the premature translation of *zelda* in embryos lacking BRAT activity, we showed that early translation of a zygotic genome activator is not sufficient to drive precocious gene expression. Instead, Zelda-target genes showed increased expression at the time they are normally activated. We propose that transition through early development requires the integration of multiple processes, including the slowing of the nuclear division cycle and activation of the zygotic genome. These processes are coordinately controlled by PNG kinase-mediated regulation of translation.

## Introduction

Regulated gene expression allows a single-celled zygote to divide and differentiate into all the cell types of the adult organism. Transcription factors bind in a sequence-specific manner to the DNA genome to drive these changes in gene expression. Nonetheless, the packaging of the genome into chromatin can limit access to the DNA, with nucleosomes acting as a barrier to transcription-factor binding (Zaret 2020). A specialized subclass of transcription factors, called pioneer factors, can bind to DNA in nucleosomes. This functionality allows pioneer factors to access regions of the genome that are inaccessible to other transcription factors. These pioneer factors facilitate chromatin opening, enabling other factors to bind and activate gene expression (Zaret 2020). Because of these unique characteristics, pioneer factors are instrumental in cellular reprogramming, and misexpression can be detrimental to the animal (Larson *et al*. 2021b). Thus, expression of pioneer factors must be precisely controlled during development.

In all animals studied to date, pioneer factors are essential for reprogramming differentiated germ cells to the totipotent embryo following fertilization (Schulz and Harrison 2019). This early developmental transition is initially controlled by maternally provided mRNAs and proteins deposited into the embryo. During this time, the zygotic genome is transcriptionally silent. Gradual activation of the zygotic genome is coordinated with the degradation of maternally deposited products during the maternal-to-zygotic transition (MZT) (Vastenhouw *et al*. 2019). Activation of zygotic transcription requires maternally encoded pioneer factors to reprogram the genome. Since the initial identification of Zelda (ZLD) as the major activator of the zygotic genome in *Drosophila melanogaster* (Liang *et al*. 2008), pioneer factors have been similarly found to be required to activate zygotic transcription in frogs, zebrafish, mice and humans (Schulz and Harrison 2019). Expression of these factors in tissues apart from the early embryo can lead to defects in development and cancer (Dobersch *et al*. 2019; Larson *et al*. 2021b).

Early *Drosophila* development is characterized by a series of 13 rapid, synchronous nuclear divisions with durations on the order of minutes. The rapidity of these divisions is ensured by the absence of any gap phases. Each division is comprised of a DNA synthesis (S phase) followed immediately by mitosis. The division cycle slows at the 14^th^ nuclear division (NC14), and this coincides with the widespread activation of transcription from the zygotic genome. However, zygotic transcription is activated in a gradual process that initiates around the 8th nuclear division (NC8) (Schulz and Harrison 2019; Vastenhouw *et al*. 2019). This activation is coordinated with the degradation of the maternally provided mRNAs that control the initial stages of development. Thus, early development requires the precise coordination of multiple processes: regulation of maternal mRNA stability and translation, transcriptional activation of the zygotic genome, and slowing of the division cycle.

The pioneer factor ZLD is a major activator of the zygotic genome in *Drosophila* (Liang *et al*. 2008; Schulz *et al*. 2015; McDaniel *et al*. 2019). *zld* is maternally deposited as an mRNA and is translationally upregulated approximately one hour after egg laying (Harrison *et al*. 2011; Nien *et al*. 2011). ZLD is bound to thousands of loci as early as NC8, marking genes that will be activated during the MZT (Harrison *et al*. 2011; Nien *et al*. 2011). In the absence of maternally deposited *zld*, embryos fail to activate their genome and die at NC14, approximately three hours after fertilization (Liang *et al*. 2008; Schulz *et al*. 2015). A mutation in *zld* that results in a hyperactive version of the protein is also lethal to the embryo (Hamm *et al*. 2017). Thus, both too much and too little of this pioneer factor activity is detrimental to development. Nonetheless, the proteins that regulate translation of *zld* mRNA and thus regulate ZLD protein levels remain largely unknown. It is also unclear whether precocious translation of *zld* would result in aberrant genome activation.

In addition to being required as a maternally contributed mRNA, ZLD is also expressed and required zygotically. Embryos lacking zygotically expressed ZLD die during embryogenesis (Liang *et al*. 2008). We recently identified a function for ZLD in a neural stem cell (neuroblast) population in the developing larval brain (Larson *et al*. 2021a). ZLD is expressed in the type II neuroblasts and rapidly eliminated from the differentiated progeny. Similar to the early embryo, ZLD levels must be precisely regulated as misexpression of ZLD in the partially differentiated progeny of the type II neuroblast lineage results in extra type II neuroblasts and a tumor-like phenotype (Larson *et al*. 2021a). During asymmetric neuroblast division, the RNA-binding protein (RBP) Brain tumor (BRAT) localizes to the differentiating daughter cell and is enriched in the partially differentiated progeny following division (Bowman *et al*. 2008; Komori *et al*. 2014). BRAT is a TRIM-NHL protein that binds (A/U)UGUU(A/G/U) motifs to regulate both mRNA stability and translation in multiple cell types, including the early embryo (Loedige *et al*. 2014, 2015; Laver *et al*. 2015). In *brat*-mutant type II neuroblast lineages, neuroblast selfrenewal factors are not downregulated and the partially differentiated progeny revert to neuroblasts (Xiao *et al*. 2012; Komori *et al*. 2014, 2018). The ectopic neuroblasts overproliferate and form tumors (Betschinger *et al*. 2006). Thus, BRAT functions to shut down the factors that maintain a neuroblast-like fate, and loss of BRAT leads to brain tumors. BRAT likely regulates ZLD levels in the type II neuroblast lineage as it can bind the *zld* 3’UTR *in vitro* and *in vivo* knockdown of *zld* can partially suppress the *brat*-mutant phenotype (Reichardt *et al*. 2018).

BRAT, like other TRIM-NHL proteins, is involved in regulating mRNA translation and stability in multiple tissues, including the early embryo (Connacher and Goldstrohm 2021). In the embryo, BRAT functions with the cofactors Pumilio (PUM) and Nanos to regulate translation of target genes such as *hunchback* (Sonoda and Wharton 2001; Loedige *et al*. 2014; Arvola *et al*. 2017). Nonetheless, recent data demonstrated that BRAT can also bind RNA independently and regulate mRNA stability (Laver *et al*. 2015; Loedige *et al*. 2015). *zld* mRNA is bound by BRAT in the early embryo but is not bound by PUM, suggesting *zld* may be directly regulated by BRAT. Gene expression analysis from embryos lacking functional BRAT identified increased levels of ZLD-target genes. Thus, BRAT may function in the early embryo to suppress ZLD levels. However, whether BRAT promotes degradation of *zld* mRNA or represses *zld* translation was unclear. Given known roles in directing post-transcriptional regulation in both the embryo and the larval neuroblasts we sought to investigate whether translational regulation of *zld* by BRAT was a shared mechanism in both tissues to limit function of this pioneer factor and to determine whether premature expression of ZLD could drive precocious activation of the zygotic genome.

## Materials and Methods

### Drosophila *strains and genetics*

All stocks were grown on molasses food at 25°C. Fly strains used in this study: *w^1118^*, *twe^HB5^ cn bw/CyO* (Courtot *et al*. 1992), *Elav-Gal4 hs-flp*, *UAS-mCD8::GFP* (BDSC#5146), *tub-Gal80, FRT40A* (BDSC#5192), *FRT40A* (Lee *et al*. 2000), *Wor-Gal4; tub-Gal80^ts^* (Komori *et al*. 2014), *y png^[50]^ w/FM6* (Fenger *et al*. 2000), *plu^[1]^/CyO* (Shamanski and Orr-Weaver 1991), *plu^[3]^/CyO* (Shamanski and Orr-Weaver 1991), *brat^150^*, *FRT40A* (Betschinger *et al*. 2006)*, bratD^f(2L)TE37C-7^/CyO* (Stathakis *et al*. 1995)*, brat^fs1^/CyO* (Schüpbach and Wieschaus 1991), and *His2Av-RFP(III)* (BDSC#23650).

*brat^150^* clones were generated by crossing *Elav-Gal4 hs-flp*, *UAS-mCD8::GFP; tub-Gal80*, *FRT40A* females to *brat^150^, FRT40A/ CyO, Act-GFP* males. *Elav-Gal4 hs-flp*, *UAS-mCD8::GFP/+* or Y; *tub-Gal80*, *FRT40A/ brat^150^*, *FRT40A* larvae were identified by lack of Act-GFP at 24 hours after egg laying. Larvae were cultured at 25°C after larval hatching. At 24 hours after larval hatching, larvae were heat shocked at 37°C for 90 min to induce clones. After the heat shock, they were cultured at 25°C again. Brains were dissected at 96 hours after larval hatching for clone analysis.

*twe^HB5^ cn bw/CyO* sterile males were crossed to *w^1118^* females to collect unfertilized embryos for western blot. *png* mutant embryos were collected from *png^[50]^*/*png^[50]^* mothers crossed to their brothers. *plu* mutant embryos were generated by crossing *plu^[1]^/CyO* and *plu^[3]^/CyO* flies. *plu^[1]^/plu^[3]^* female progeny were then crossed to their brothers, and the embryos from these *plu^[1]^/ plu^[3]^* mothers were collected for western blot. *brat-*mutant embryos were generated by crossing *brat^Df(2L)TE37C-7^/CyO* and *brat^fs1^/CyO. brat^Df(2L)TE37C-7/fs1^* female progeny were collected and crossed to their brothers. The embryos from these *brat^Df(2L)TE37C-7/fs1^* mothers were collected for western blot and qRT-PCR. *brat^Df(2L)TE37C-7^/CyO;His2Av-RFP(III)* (generated for this manuscript) were used in crosses to generate *brat-*mutant embryos for RNA-seq and nuclear-cycle timing. The *brat^fs1^* mutation results in a G774D amino acid substitution in the NHL domain that causes female sterility (Schüpbach and Wieschaus 1991; Arama *et al*. 2000; Sonoda and Wharton 2001).

We initially planned to use embryos that were heterozygous for the *brat*-mutant alleles (*brat* hets) as controls (*brat^fs1/+^* and *brat^Df/+^*). However, upon further analysis we discovered that these heterozygous embryos display differential gene expression as compared to *His2AvRFP* embryos (Supplemental Figure 1). At NC10 over half of the genes that were mis-regulated in the *brat*-mutant embryos were also mis-regulated in the embryos laid by *brat* hets (Supplemental Figure 1A). Not one particular allele contributes to these changes as when we overlapped the genes that were mis-regulated in the *brat^fs1/+^* and *brat^Df/+^* NC10 embryos, a large percentage of them are shared (Supplemental Figure 1C,D). Additionally, we found that many genes were already mis-regulated in a *brat*-mutant oocyte. And many of these mis-regulated genes were similarly mis-regulated in the *brat* het oocytes (Supplemental Figure 1B). Based on the observed gene expression changes in the embryos laid by heterozygotes, we decided to use *His2AvRFP* flies (*HisRFP*) as controls.

**Figure 1:**
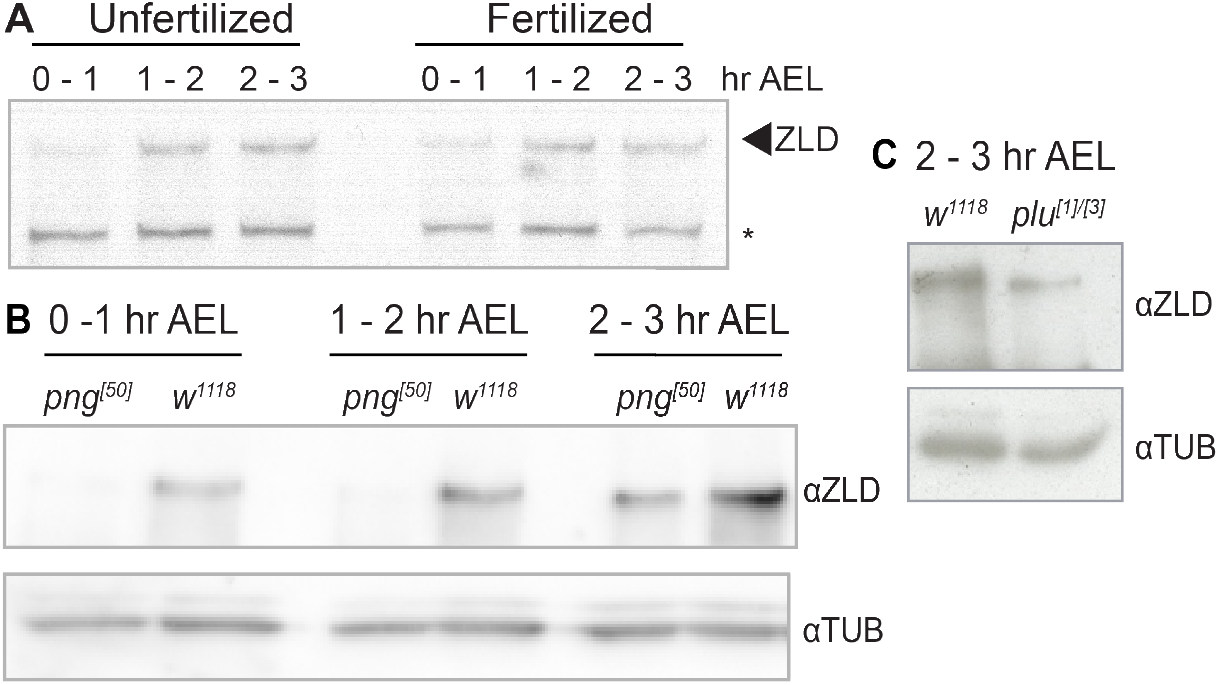
Translational upregulation of ZLD is reduced in PNG kinase complex mutants. A. Immunoblots with anti-ZLD antibody on unfertilized or fertilized embryos collected at one-hour time points after egg laying (AEL). Non-specific background (*) band serves as loading control. B. Immunoblots on control (*w^1118^*) or *png*-mutant embryos at one-hour intervals after egg laying (AEL). Tubulin is shown as a loading control. C. Immunoblots on control (*w^1118^*) or *plu-mutant* embryos at 2-3 hours after egg laying (AEL). Tubulin is shown as a loading control.

To generate transgenes for ectopic expression of ZLD regulated by various 3’UTRs, we cloned the open reading frame for ZLD-RB and 3’UTRs into pUASt-attB using standard PCR, restriction digest and ligation procedures. BRE mutations in the 3’UTR of *zld* were cloned using iPCR and gBlocks containing BRE mutations (Integrated DNA Technologies, Coralville, IA). BREs were mutated from UGUU to CGCU. These transgenes were integrated at ZH-86Fb site on chromosome 3 using φC31-mediated integration (Bischof and Basler 2008) (BestGene, Chino Hills, CA). The *Wor-Gal4; tub-Gal80^ts^* driver was used to drive ectopic expression of ZLD in the larval type II neuroblast.

### Immunoblotting

Proteins were transferred to 0.45 μm Immobilon-P PVDF membrane (Millipore, Burlington, MA) in transfer buffer (25 mM Tris, 200 mM Glycine, 20% methanol) for 75 min at 500mA at 4°C. The membranes were blocked with blotto (2.5% non-fat dry milk, 0.5% BSA, 0.5% NP-40, in TBST) for 30 min at room temperature and then incubated with rabbit anti-ZLD (1:750) (Harrison *et al*. 2010), or anti-Tubulin (DM1A, 1:5000) (Sigma, St. Louis, MO), overnight at 4°C. The secondary incubation was performed with goat anti-rabbit IgG-HRP conjugate (1:3000) (Bio-Rad, Hercules, CA) or anti-mouse IgG-HRP conjugate (1:3000) (Bio-Rad) for 1 hr at room temperature. Blots were treated with SuperSignal West Pico PLUS chemiluminescent substrate (Thermo Fisher Scientific, Waltham, MA) and visualized using the Azure Biosystems c600 or Kodak/Carestream BioMax Film (VWR, Radnor, PA).

### Immunofluorescent staining of larval brains and antibodies

Third instar larval brains were dissected in PBS and fixed in 100 mM Pipes (pH 6.9), 1 mM EGTA, 0.3% Triton X-100 and 1 mM MgSO4 containing 4% formaldehyde for 23 minutes. Fixed brain samples were washed with PBST containing PBS and 0.3% Triton X-100. After removing fix solution, samples were incubated with primary antibodies for 3 hours at room temperature. Samples were washed with PBST and incubated with secondary antibodies for overnight at 4°C. The next day samples were washed with PBST and then equilibrated in ProLong Gold antifade mountant (ThermoFisher Scientific, Waltham, MA). The confocal images were acquired on a Leica SP5 scanning confocal microscope (Leica Microsystems Inc, Buffalo Grove, IL). 10 brains per genotype were used to obtain data in each experiment. Antibodies used include rabbit anti-ZLD (1:500) (Harrison *et al*. 2010), rat anti-DPN Antibody (1:2) (clone 11D1BC7.14) (Lee *et al*. 2006), rabbit anti-ASE (1:400) (Weng *et al*. 2010) and chicken anti-GFP Antibody (1:2000) (Aves Labs, Davis, CA, Cat #GFP-1020).

### *Quantification of* zld *mRNA levels in* brat-*mutant embryos*

To assess *zld* transcript levels in wild-type and *brat*-mutant embryos, total RNA was isolated (see RNA-seq) from 10 embryos collected from *w^1118^* or *brat*-mutant females (as described above). The total RNA was then used to prepare single-stranded cDNA by reverse transcription using random primers and Superscript IV reverse transcriptase (Invitrogen, Waltham, MA). The single-stranded cDNA was used to perform quantitative real-time PCR with primers specific to the *zld* transcript, as well as *Act5C* (see reagents table for primer sequence) as a control mRNA that is unaffected in *brat* mutants, using GoTaq qPCR Master Mix (Promega, Madison, WI, Cat #A6001).

### Single oocyte and embryo RNA-seq

Single *bratD^f/fs1^;His2Av-RFP(III)*, *brat het;His2Av-RFP(III)*, or *His2Av-RFP(III)* embryos (3-4 replicates per time point) were dechorionated in 100% bleach for 1’, mounted in halocarbon (700) oil (Sigma Aldrich, St. Louis, MO) on a coverslip, and imaged using on a Nikon Ti-2e Epiflourescent microscope using a 60X objective. The nuclear cycle was identified following mitosis based on His2Av-RFP marked nuclear density (calculated by the number of nuclei/2500 μm^2^). At the indicated time, embryos were picked into Trizol (Invitrogen, Waltham, MA, Cat #15596026) with 200 μg/ml glycogen (Invitrogen, Waltham, MA, Cat #10814010) and pierced with a 27G needle to release the RNA for 5 min. Late-stage single oocytes (4 replicates per genotype) were dissected from the ovaries of the mothers of the respective genotype and staged based on morphology. The oocytes were picked into Trizol (Invitrogen, Waltham, MA, Cat #15596026) with 200 μg/ml glycogen (Invitrogen, Waltham, MA, Cat #10814010) and pierced with a 27G needle to release the RNA for 5 min. RNA was extracted and RNA-seq libraries were prepared using the TruSeq RNA sample prep kit v2 (Illumina, San Diego, CA). 75-bp reads were obtained using an Illumina NextSeq500 sequencer at Northwestern Sequencing Core (NUSeq Core). This protocol is expanded in McDaniel and Harrison 2019.

### RNA-seq analysis

Raw reads were aligned to the BDGP *D. melanogaster* genome release 6 (dm6) using hisat2 v2.1.0 with the following parameters: -k 2 –very-sensitive (Kim *et al*. 2015). Reads aligning to multiple locations were discarded. SAMtools was used to filter (-q 30) and convert file formats (Li *et al*. 2009). Reads were assigned to annotated genes using featureCounts v1.5.3 (Liao *et al*. 2014) using default parameters and the UCSC annotation (r6.20). The resultant table of read counts was imported into R v4.1.0, and differential expression was determined using DESeq2 v1.33.5. (Love *et al*. 2014). Genes with < 50 reads across all samples were filtered out. Differential expression (adjusted p value < 0.05 and log_2_(fold change) > 1) was determined using the standard DESeq2 analysis. Read counts were z-score normalized for visualization of the time course line graphs. To identify ZLD-target genes, previously determined ZLD-ChIP peaks were assigned to the nearest gene (ChIP from Harrison *et al*. 2011, reanalyzed by Larson *et al*. 2021a). Previously identified BRAT-regulated transcripts and BRAT-bound transcripts were from Laver *et al*. 2015. Zygotically and maternally expressed genes, as well as the timing of zygotic gene expression, were previously defined (Lott *et al*. 2011; Li *et al*. 2014; Strong *et al*. 2020). Lists of differentially expressed genes are in Supplementary Table 1.

### Nuclear-cycle timing

Single *brat^Df(2L)TE37C-7/fs1^;His2Av-RFP(III)* embryos were prepared and mounted as for RNA-seq above. S-phase was timed from the exit of mitosis to the first sign of nuclei condensation.

## Results

### ZLD levels depend on the Pan Gu kinase

*zld* mRNA is deposited in the embryo and translated approximately one hour after fertilization (Harrison *et al*. 2010; Nien *et al*. 2011). To understand how ZLD protein levels are precisely controlled, we determined whether the timing of *zld* translation depended on fertilization of the oocyte. Immunoblots on both fertilized and unfertilized embryos harvested in one-hour increments after egg laying (AEL) showed that ZLD protein levels increased regardless of whether the oocytes were fertilized (Figure 1A). Thus, a maternally regulated process controls the timing of *zld* translation.

Translation in the early embryo is broadly regulated by the Pan Gu (PNG) kinase, which phosphorylates and inactivates translational repressors (Kronja *et al*. 2014a, 2014b; Eichhorn *et al*. 2016; Hara *et al*. 2018). PNG is comprised of three subunits (Pangu (PNG), Giant nuclei (GNU) and Plutonium (PLU)) and is subject to precise control such that it is active in a short window following egg activation (Hara *et al*. 2017). Embryos with mutations in PNG subunits arrest in early development with failures in nuclear division resulting from the failure to translate *cyclin B* (Shamanski and Orr-Weaver 1991; Fenger *et al*. 2000; Lee *et al*. 2003; Vardy and Orr-Weaver 2007). We performed immunoblots for ZLD in embryos lacking functional PNG subunits, PNG or PLU. ZLD levels were reduced in embryos laid by *png^[50]^* mutant mothers during the three hours AEL as compared to *w^1118^* control embryos (Figure 1B). Similarly, in 2-3 hr AEL embryos laid by *plu^[1]/[3]^* mutant mothers, ZLD levels were lower than controls (Figure 1C). Because ZLD translation occurs in the absence of fertilization, we can conclude that the decrease in ZLD levels in these mutants is due to defects in PNG kinase activity rather than the failure of the embryos to develop.

### BRAT regulates ZLD levels in larval neuroblasts through consensus binding sites

As in the early embryo, ZLD levels in the type II neuroblast lineage of the larval brain are tightly controlled (Reichardt *et al*. 2018; Larson *et al*. 2021a). In the neuroblast lineage, ZLD levels are controlled by Brain Tumor (BRAT), which binds to *zld* mRNA and has been suggested to degrade it (Reichardt *et al*. 2018). In wild-type animals, ZLD is rapidly eliminated from the partially differentiated progeny (immature intermediate neuroblast progenitor (immINP)) following asymmetric neuroblast division (Figure 2A,B) (Reichardt *et al*. 2018). By contrast, in a *brat* mutant, ZLD protein is retained in both cells following division (Figure 2B) (Reichardt *et al*. 2018). The *zld* 3’UTR contains multiple BRAT-binding elements (BREs) (Figure 2C), and these can be bound by BRAT *in vitro* (Reichardt *et al*. 2018).

**Figure 2:**
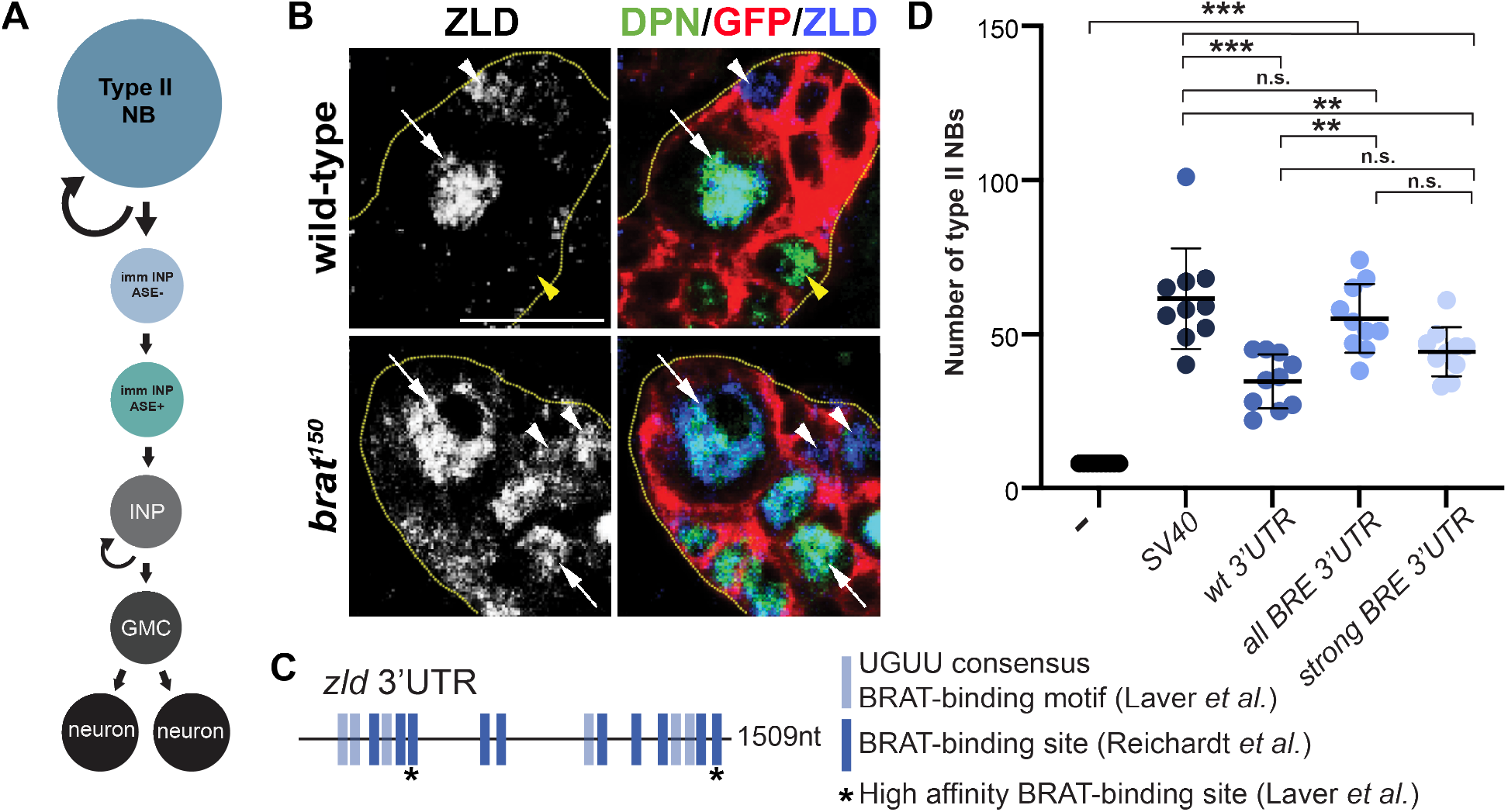
BRAT-binding sites in the 3’UTR of *zld* regulate ZLD activity in the larval neuroblasts. A. Schematic of the type II neuroblast (NB) lineage. immlNP = immature neural progenitor (INP). GMC = ganglion mother cell. B. Immunostaining of wild-type or *brat^150^* clones in the third instar larval brains for Deadpan (DPN) (a marker of neuroblasts and INPs), ZLD and GFP to mark the clone. Dashed yellow lines mark the clone boundary. White arrows indicate type II neuroblasts. White arrowheads represent Asense negative (ASE-) immature INPs, and yellow arrowheads indicate mature INPs. Scale bar = 10mm. C. Schematic of the *zld* 3’UTR with previously identified BRAT (UGUU) motifs highlighted. Asterisks indicate two strong binding sites as determined by RNAcompete. D. Quantification of type II neuroblasts when ZLD is overexpressed from reporters with the 3’UTRs indicated below. Error bars indicate standard deviation. ** = p < 0.005, *** = p < 0.0001 and n.s. = non-significant as determined by a one-way ANOVA test with post hoc Tukey’s multiple comparisons test.

In the larval brain there are exactly eight type II neuroblast per lobe. Loss-of-function in *brat* results in extra neuroblasts, and this can be partially suppressed by knockdown of *zld* (Reichardt *et al*. 2018). We recently demonstrated that in wild-type brains overexpression of *zld* with the SV40 3’UTR, which is commonly used for transgene expression and stabilizes mRNA, results in extra stem cells (61.5±15.9 type II neuroblasts) (Figure 2D) (Larson *et al*. 2021a). We therefore used this quantitative system to determine whether BRAT mediates regulation of ZLD levels through binding to the 3’UTR. Expression of *zld* with the endogenous 3’UTR resulted in fewer extra neuroblasts (34.7±8.5 type II neuroblasts) as compared to *zld* with the SV40 3’UTR (Figure 2D), supporting a role for the 3’UTR in regulating ZLD activity in the neuroblasts. We then mutated all 16 identified BREs in the *zld* 3’UTR. Overexpression of *zld* regulated by the 3’UTR containing mutations in all 16 putative BREs resulted in significantly more neuroblasts (55.1±10.9 type II neuroblasts) than the wild-type *zld* 3’UTR (p-value = 0.0006, ANOVA with post hoc Tukey’s multiple comparisons test) (Figure 2D). RNAcompete predicted that two of the BREs in the *zld* 3’UTR were high-affinity (strong) BREs (Figure 2A) (Laver *et al*. 2015). To test the relative contribution of these two BREs, we created transgenes in which only these two BREs were mutated in the *zld* 3’UTR. When compared to the number of neuroblasts induced when all 16 BREs were mutated, mutation of only the strong BREs did not result in a significant difference (44.3±7.8 type II neuroblasts). Together these data support a role for BRAT in regulating ZLD levels in the neuroblast lineage by binding preferentially to a subset of BREs in the 3’UTR.

### BRAT regulates ZLD levels in the early embryo

In the early embryo, BRAT regulates both mRNA translation and stability and is bound to *zld* mRNA (Laver *et al*. 2015). This, together with our demonstration of the role of BRAT in regulating *zld* in the neuroblasts, suggested that BRAT may regulate *zld* translation in the early embryo. To test this, we immunoblotted for ZLD in staged control (*w^1118^*) and *brat*-mutant embryos (embryos laid by *brat^Df/fs1^* mothers) (Figure 3A). *brat*-mutant embryos are devoid of functional BRAT and do not survive past embryogenesis (Schüpbach and Wieschaus 1991; Stathakis *et al*. 1995; Arama *et al*. 2000; Sonoda and Wharton 2001; Loedige *et al*. 2014). In *w^1118^* embryos ZLD levels increased normally from 0-3 hr AEL. By contrast, at 0-1 hr AEL in *brat*-mutant embryos ZLD protein levels were already at levels similar to controls at 1-2 hr AEL, after *zld* translation increased (Figure 3A). This premature increase in ZLD levels in *brat*-mutant embryos is not due to changes in the total RNA present. RT-qPCR for *zld* in control (*w^1118^*) and *brat*-mutant embryos at two time points, stage 2-3 and stage 5, showed that *zld* mRNA levels remained relatively constant as compared to an internal control, *Act5C* (Figure 3B). Together with previously published data, this suggests that BRAT binds to *zld* mRNA and represses translation immediately following fertilization.

**Figure 3:**
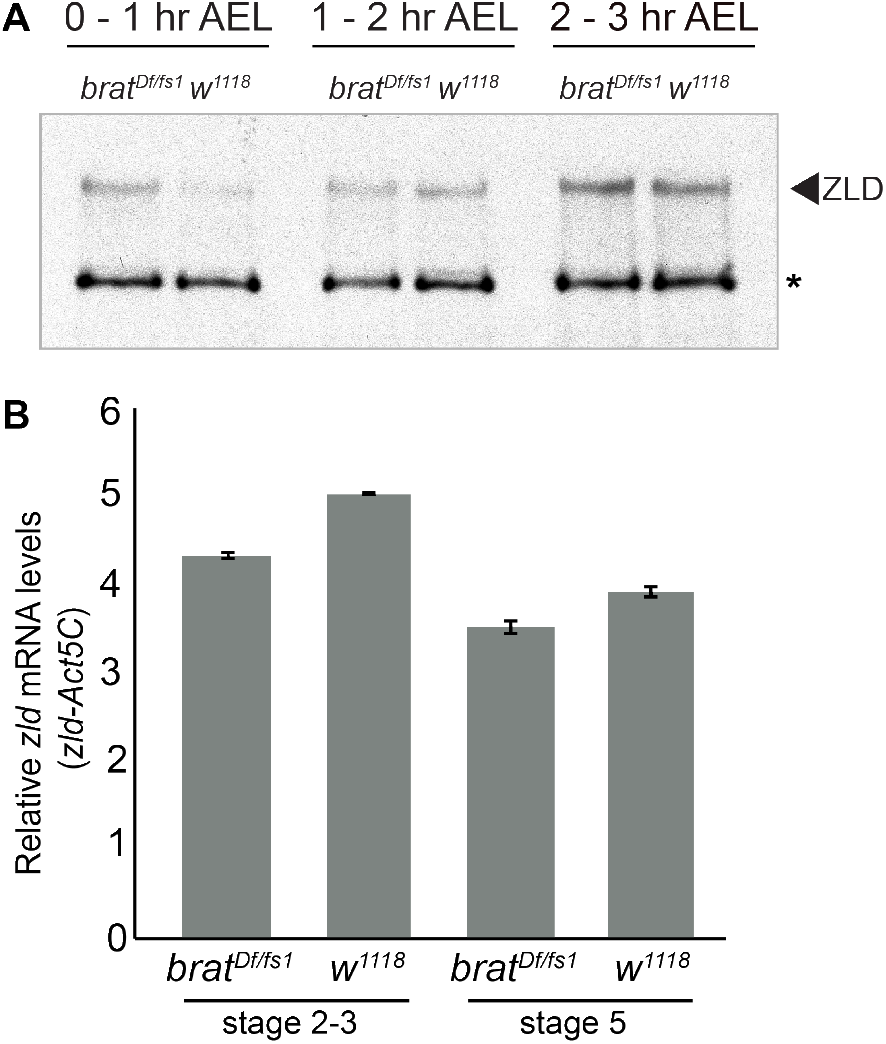
BRAT regulates ZLD levels in the early embryo. A. Immunoblots on control (*w^1118^*) or *brat^Df/fs1^* embryos at one-hour intervals after egg laying (AEL). Non-specific background band (*) serves as a loading control. B. RT-qPCR quantification of *zld* mRNA levels as compared to *Act5C* control. Error bars indicate normalized standard deviation from two biological replicates.

### Premature ZLD expression results in increased, but not precocious target-gene expression

Our data suggested that translational repression of *zld* by BRAT immediately following fertilization was alleviated, directly or indirectly, by the activity of the PNG kinase and prior data demonstrated that too much ZLD activity was lethal to the embryo. Based on these observations, we hypothesized that precocious translation of ZLD might be detrimental to the embryo by driving premature expression of ZLD-target genes. Supporting this hypothesis, these ZLD-target genes had increased expression in *brat*-mutant embryos 1.5-3 hr AEL as assayed by microarray (Laver *et al*.2015). At this early stage of embryonic development, the syncytial nuclei are rapidly dividing, and transcription is gradually activated over about one hour of development, culminating in a major wave of gene expression when the nuclear division cycle slows at approximately two hours after fertilization. Thus, based on the published microarray data that was generated from bulk collections of timed embryos, it was not possible to determine whether the identified increase in ZLD-target gene expression was due to precocious gene expression or, instead, an increase in expression at the time of normal transcriptional activation.

To determine whether premature ZLD expression can drive precocious gene expression, we performed single-embryo RNA-seq on *brat*-mutant and control (*HisRFP*) embryos at six precise time points spanning late oocytes and early embryogenesis. Embryos were precisely staged based on imaging the nuclear density of a fluorescently tagged histone (His2Av-RFP). RNA-seq was performed on single oocytes and single embryos at NC10, half-way into NC12 (NC12^+5min^), half-way into NC13 (NC13^+8min^), at the beginning of NC14 (NC14^+5min^) and half-way into NC14 (NC14^+30min^) (Figure 4A). To validate the data collected, we performed principal component analysis (PCA). We included previously published single-embryo RNA-seq data from wild-type embryos for comparison (Lott *et al*. 2011). The PCA demonstrated that the oocyte and embryo replicates collected at each time point cluster together and cluster with independently collected, previously published data from wild-type embryos (Figure 4B). Further supporting the robustness of the data collected, the replicates from each time point of each genotype were highly correlated (Supplemental Figure 2).

**Figure 4:**
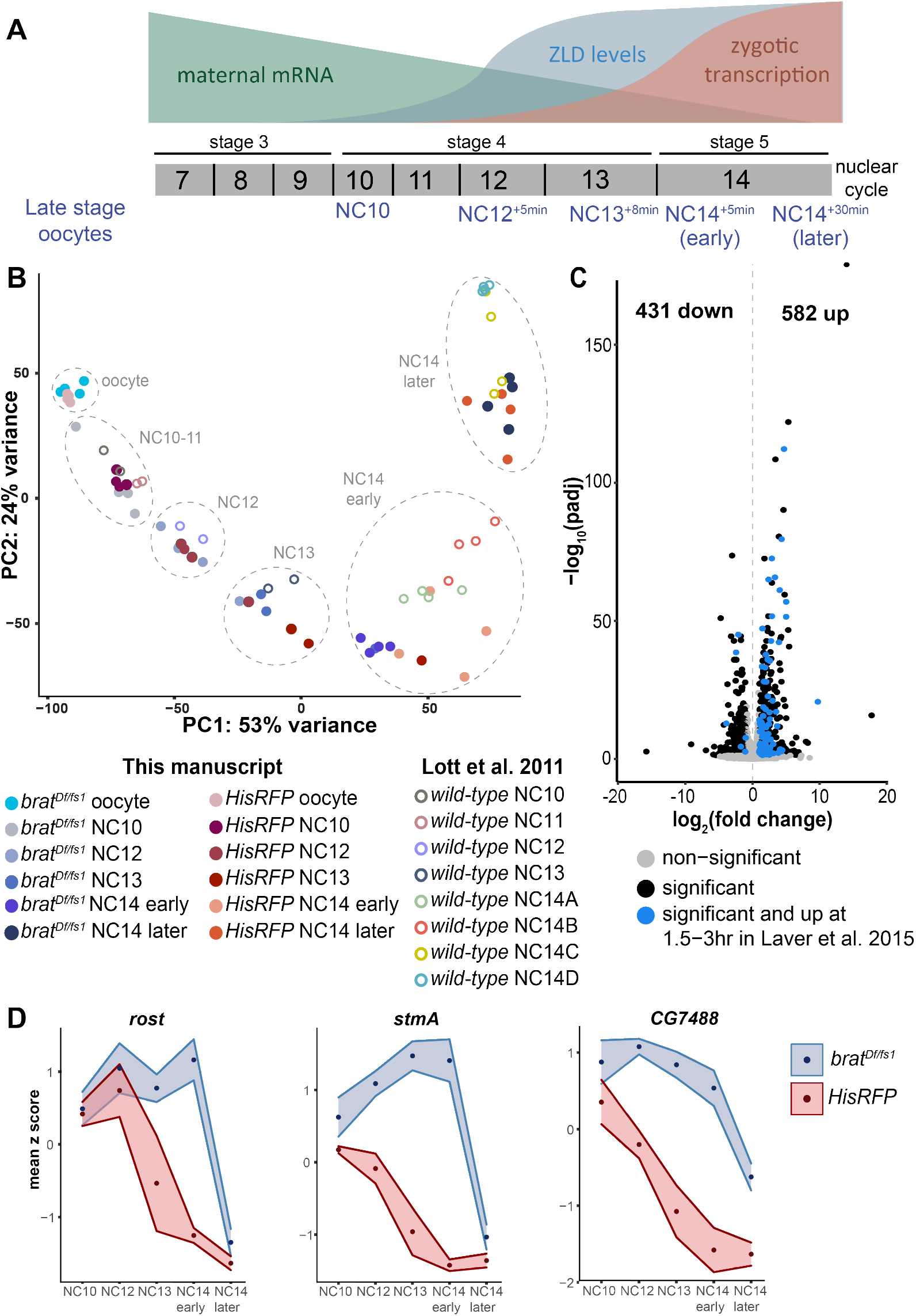
Identification of genes mis-regulated in *brat*-mutant embryos. A. Developmental timing of oocytes and embryos collected for RNA-seq. B. PCA plot of RNA-seq from embryos/oocytes. Previously published single-embryo, RNA-seq data from wild-type embryos are included as verification of developmental staging (Lott *et al*. 2011). C. Volcano plot highlighting genes mis-regulated at NC14 in *bratD^f/fs1^* embryos as compared to *HisRFP* controls. Blue dots indicate genes that are were also increased in *brat* mutants at 1.5 - 3 hr AEL (Laver *et al*. 2015). Black dots indicate genes not identified in Laver *et al*. 2015 that change in expression in *brat^Df/fs1^* embryos as compared to controls (log_2_(fold change) >1, padj <0.05). D. Expression pattern of three genes bound by BRAT (Laver *et al*. 2015) with increased expression in *brat^Df/fs1^* embryos as compared to controls at all developmental timepoints analyzed. Points indicate the average z score, and the surrounding region indicates the standard deviation of the replicates for each timepoint.

We identified gene expression changes in *brat*-mutant embryos as compared to control embryos at NC14^+30min^, which is most similar to the prior time point analyzed by Laver *et al*. 2015. We identified 582 genes with significantly higher mRNA signal in the *brat*-mutant embryos as compared to the controls (Figure 4C). These genes are comprised of those that were previously found to be increased in the published microarray data (Figure 4C) (Laver *et al*. 2015). Many of the genes that have increased mRNA levels are bound by BRAT in wild-type embryos and are stabilized in the *brat* mutant (Laver *et al*. 2015). These are exemplified by *rolling stone* (*rost*), *stambha A* (*stmA*) and *CG7488*, which are directly regulated by BRAT binding (Figure 4D). These analyses provided confidence in the data generated and allowed us to interrogate these data to determine if ZLD-target genes were precociously expressed.

We initially focused our analysis on genes that were increased in the *brat*-mutant embryos as compared to controls at any of the embryonic time points collected (NC10 through NC14^+30min^). We performed k-means clustering on the expression of this set of genes in the control embryos and identified six clusters with distinct expression patterns (Figure 5A). Clusters 1-4 decreased in expression over the time course. These transcripts were enriched for maternally deposited genes and most were bound by BRAT (Lott *et al*. 2011; Laver *et al*. 2015; Strong *et al*. 2020). Thus, these clusters are largely comprised of maternal genes that are direct BRAT targets and are normally degraded during the MZT. By contrast, clusters 5-6 increased in expression over the time course. The transcripts in these clusters were enriched for zygotically expressed genes and the majority were not bound by BRAT (Figure 5A) (Lott *et al*. 2011; Laver *et al*. 2015; Strong *et al*. 2020). As might be expected for zygotically expressed genes, the genes in clusters 5 and 6 are located proximally to regions occupied by ZLD in the early embryo (Harrison *et al*. 2011). Thus, these clusters contain ZLD-target genes that are indirectly increased in the *brat* mutants and therefore are likely to include any genes that might be precociously activated by the premature translation of ZLD in the *brat*-mutant embryos.

**Figure 5:**
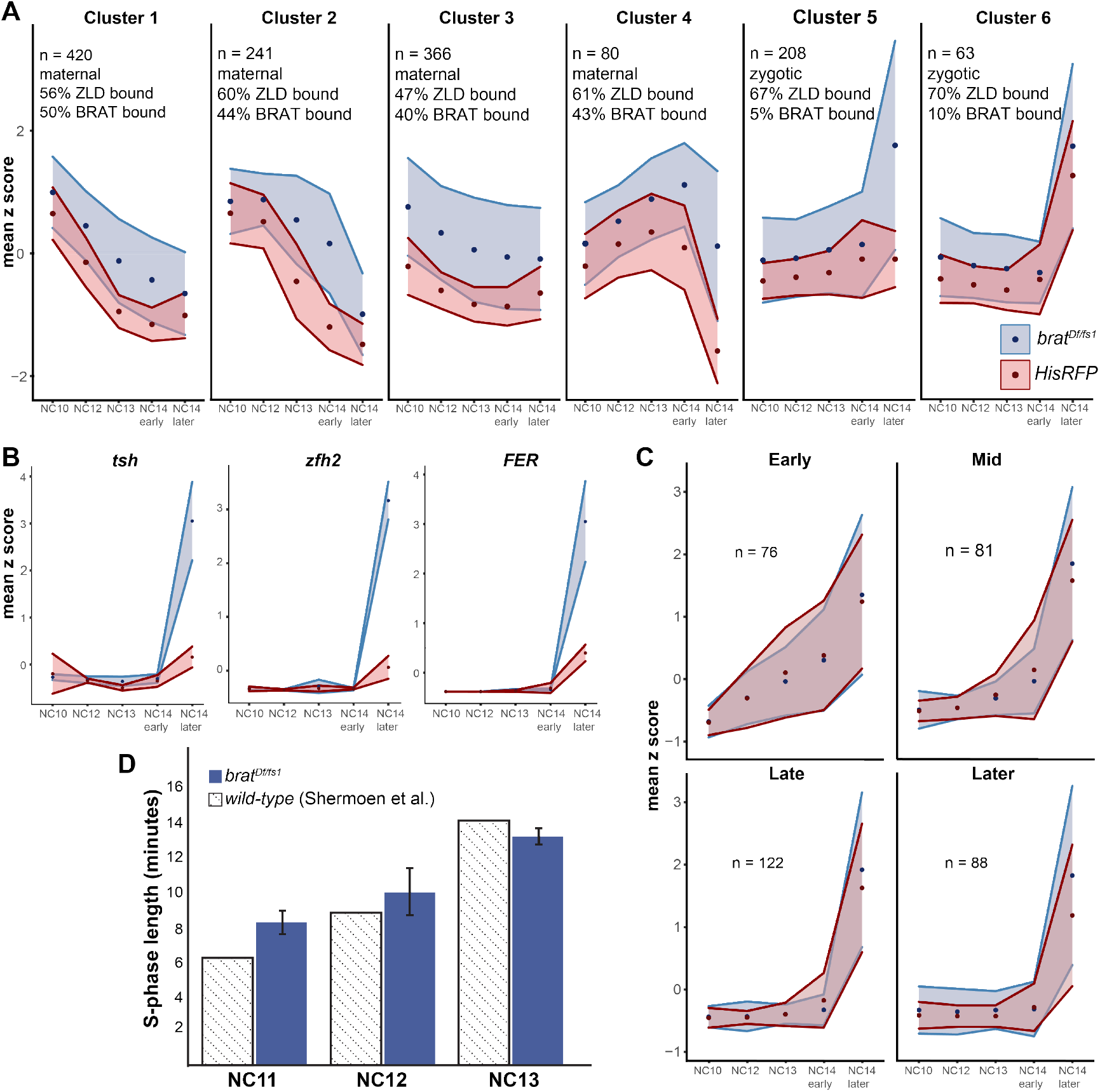
ZLD-target genes are increased in expression in *brat*-mutant embryos at NC14 but not earlier. A. k-means clustering of all genes significantly increased in *brat^Df/fs1^* embryos as compared to *HisRFP* controls (log_2_(fold change) >1, padj <0.05). Points indicate the average z score, and the surrounding region indicates the standard deviation of the replicates for each timepoint. n = number of genes in each cluster. B. Expression for three representative genes that are bound by ZLD, not bound by BRAT, and significantly increased in *brat^Df/fs1^* as compared to *HisRFP* control embryos (cluster 5). C. Expression profiles of classes of zygotically expressed genes (Li *et al*. 2014). D. Length in minutes for S phase of nuclear cycles 11-13 in *brat^Df/fs1^* embryos and wild-type embryos as determined by Shermoen *et al*. 2010. Error bars are standard deviation of 3 biological replicates.

Having identified those genes most likely to reflect increased activation due to precocious ZLD expression, we leveraged our tightly staged RNA-seq data to determine the dynamics of gene expression that generated the observed increase in transcript levels. If these genes are precociously activated, they would be expected to have increased transcript levels in the *brat* mutant as compared to controls at time points preceding widespread gene activation at NC14 (NC12^+5min^, NC13^+8min^, NC14^+5min^). In contrast to this prediction, genes in clusters 5 and 6 do not exhibit increased transcript abundance at these early time points (Figure 5A). Instead, the largest increase in expression of genes in a *brat*-mutant embryo is evident at NC14^+30min^. ZLD-target genes *teashirt (tsh), zinc finger homeodomain 2 (zfh2*) and *FER* exemplify this robust hyperactivation at NC14 (Figure 5B).

To further investigate whether a small subset of zygotically transcribed genes might be precociously activated, we classified genes based on their timing of transcriptional activation, early (NC10-11), mid (NC12-13), late (early NC14) or later (late NC14) (Li *et al*. 2014) and plotted their expression in *brat*-mutant and control embryos (Figure 5C). Genes that initiated transcription prior to the major wave at NC14 (early, mid, and late) showed relatively little difference in expression levels between the control and *brat* mutant (Figure 5C). By contrast for the genes activated later in NC14, transcript levels were increased in the *brat*-mutant embryos as compared to controls (Figure 5C). Similar to what we observed with our k-means clustering analysis, zygotic ZLD-target genes were not precociously activated in the *brat*-mutant embryos. Instead, an increased level of zygotic transcripts was evident at NC14. We conclude that the premature increase in ZLD protein in *brat*-mutant embryos results in increased, rather than precocious, expression of zygotic ZLD-target genes.

Our data demonstrate that precocious expression of a major activator of the zygotic genome is not sufficient to drive gene expression early. Thus, additional factors are required for timing the transcriptional activation of the zygotic genome. During early development, the nuclei are dividing rapidly (~every 10 minutes). It is not until NC14 that this division cycle slows. Because the division cycle is known to disrupt transcription, we hypothesized that these rapid divisions may keep precocious ZLD from activating transcription globally until NC14. Indeed, we timed the length of S phase in *brat*-mutant embryos at NC11, NC12 and NC13 and confirmed that the nuclear division cycles occurred in these mutants with the same timing as wild-type (Figure 5D) (Shermoen *et al*.2010). Together our data demonstrate that precocious expression of an essential activator of the zygotic genome is not capable of driving early gene expression, and we suggest that the slowing of the cell cycle is essential for the activator to promote robust gene expression.

## Discussion

Here we demonstrate that BRAT regulates levels of the pioneer transcription factor ZLD in both the larval neuroblasts and in the early embryo. In both tissues, this regulation is required for normal development. In the neuroblasts, BRAT functions to ensure that ZLD levels are rapidly reduced following asymmetric division, enabling cells to exit the stem-cell fate and begin to differentiate. We previously demonstrated that following asymmetric neuroblast division misexpression of ZLD in the differentiating progeny results in a reversion to the neuroblast fate and a tumor-like phenotype (Larson *et al*. 2021a). Furthermore, knockdown of *zld* can partially rescue a *brat*-mutant phenotype in the larval brain (Reichardt *et al*. 2018). Our data from transgenes in which *zld* is expressed with wild-type or mutated 3’UTRs show that BRAT-binding sites in the 3’UTR function to suppress ZLD activity following asymmetric division. BRAT is localized to the differentiating progeny and promotes deadenylation and subsequent degradation of the target gene *deadpan* (Komori *et al*. 2014; Reichardt *et al*. 2018). Thus, BRAT may similarly promote degradation of *zld* mRNA in the differentiating stem-cell progeny, resulting in decreased ZLD protein.

Normal development also requires that ZLD levels be precisely controlled in the early embryo. At this time in development, the genome is transcriptionally quiescent. Therefore, protein levels are controlled via regulation of mRNA stability and translation. ZLD is encoded by a maternally provided mRNA but is not translated until approximately 1 hour after fertilization. We show that BRAT represses *zld* translation immediately following fertilization and that this repression is alleviated by the activity of the PNG kinase (Figure 6). *zld* RNA levels do not increase in *brat*-mutant embryos, suggesting that BRAT regulates ZLD protein levels by repressing translation. This mechanism may differ from BRAT-mediated regulation of *zld* in the neuroblast lineage. It has been previously shown that BRAT can post-transcriptionally regulate transcripts through either RNA degradation or translational repression so it is possible that BRAT regulates ZLD levels differently in different tissues (Connacher and Goldstrohm 2021). Indeed, the early embryo is unique in that it is the only developmental time point at which poly(A)-tail length is correlated with translation efficiency (Eichhorn *et al*. 2016). Thus, BRAT-mediated deadenylation could result in translational repression rather than degradation at this early time in development. However, BRAT also more directly regulates translation initiation through interactions with the cap binding protein, 4EHP (Connacher and Goldstrohm 2021). In addition to the role of BRAT in repressing *zld* translation, BRAT is also required during the MZT for degradation of maternally provided mRNAs (Laver *et al*. 2015). Thus, the function of BRAT may depend both on the target mRNAs bound and developmental stage. Indeed, this duality is not limited to BRAT as ME31B represses translation early during the MZT, but transitions to promoting mRNA destruction as development proceeds in a manner regulated by PNG (Wang *et al*. 2017).

**Figure 6:**
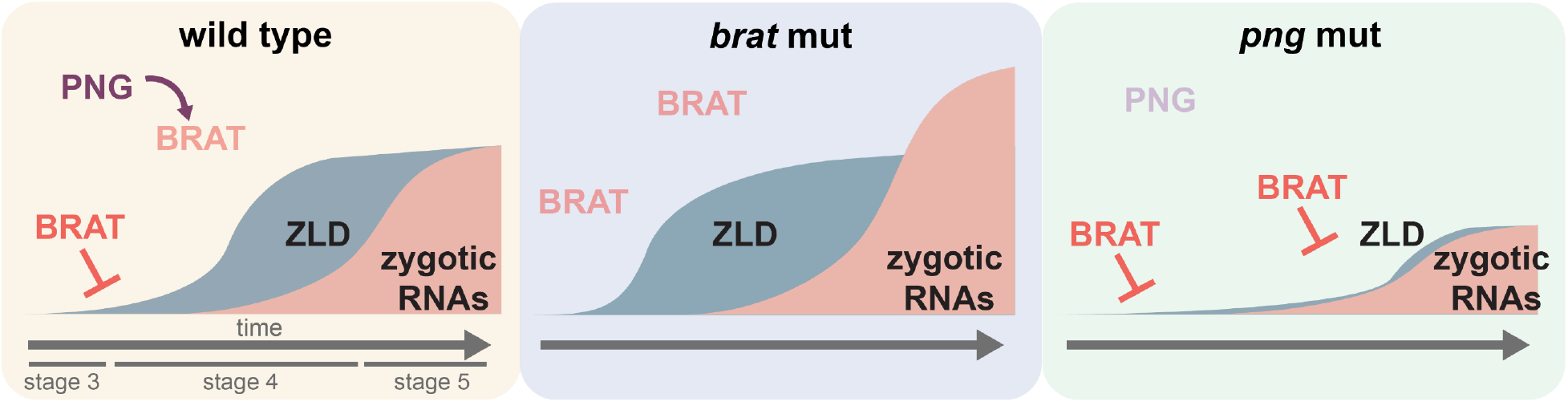
PNG kinase activity alleviates translational repression of *zld* by BRAT. In wild-type embryos BRAT represses *zld* translation during the earliest stages of embryogenesis, and this repression is alleviated by the PNG kinase complex. In a *brat*-mutant embryo, *zld* is precociously translated. This results in increased transcription of ZLD-target genes at stage 5, the time in which they are normally expressed. When PNG activity is lost, *zld* translation is inhibited throughout the MZT, and ZLD expression fails to reach wild-type levels. We propose PNG activation coordinates multiple essential processes during the MZT including increased levels of the zygotic genome activator, ZLD.

Having demonstrated precocious expression of ZLD in *brat*-mutant embryos, we used this system to test whether early expression of a zygotic genome activator could prematurely drive transcriptional activation. Using RNA-seq on precisely staged embryos, we showed that while transcript levels of ZLD-target genes were increased in the *brat* mutant this occurred only at the time at which these genes would normally be activated.

Early models suggested that the ratio of nuclear content to cytoplasmic content was essential for genome activation and that titration of a maternally provided repressor was required for genome activation (Newport and Kirschner 1982; Schulz and Harrison 2019). Work from zebrafish and *Xenopus* identified histones as maternally provided repressors and showed that increased concentrations of pioneer factors allow these genome activators to outcompete histones to activate gene expression (Amodeo *et al*. 2015; Joseph *et al*. 2017). In contrast to these models, our data indicate that increased concentration of pioneer factors alone is not sufficient to precociously activate zygotic transcription. Instead, developmental features aside from the increased levels of the transcriptional activators are required for widespread activation of the zygotic genome. We propose that one of these features is the slowing of the division cycle. As in many other organisms, the initial division cycles in the *Drosophila* embryo are extremely rapid, only lasting about 10 minutes. During these rapid cycles, transcription is limited to S-phase as mitosis interrupts transcription (McCleland *et al*. 2009). It is not until NC14 that the division cycle slows and provides ample time for robust gene expression. Further supporting a process independent from nuclear to cytoplasmic ratio in timing activation of the zygotic genome, recent studies in *Drosophila* and zebrafish showed that zygotic genome activation of a large subset of genes can occur in the absence of increased nuclear content (Chan *et al*. 2019; Strong *et al*. 2020). Instead, this activation requires both translation and time after fertilization (Chan *et al*. 2019; Strong *et al*. 2020). In zebrafish, as in flies, the levels of the major activators of the zygotic genome Pou5f3, Sox19b, and Nanog are controlled by robust translation following fertilization (Lee *et al*.2013). Together with our data, this suggests that translation of genome activators coupled with a slowing of the division cycle is essential for zygotic genome activation.

Our data support a model in which activation of the zygotic genome requires a tight coordination of multiple processes, including increased levels of zygotic genome activators and slowing of the division cycle. We propose that in *Drosophila* activation of the PNG kinase coordinately regulates these processes and couples them to egg activation, allowing for robust control of this process. PNG kinase is essential for translation of *cyclin B*, which encodes a key regulator of the division cycle in the early embryo, and *smaug*, which regulates maternal mRNA stability (Tadros *et al*. 2007; Vardy and Orr-Weaver 2007). Here we show that *zld* translation is similarly dependent on PNG activity (Figure 6). PNG phosphorylates key regulators of translation, ME31B, BIC-C and Trailer hitch, and, in the case of TRAL, blocks it’s repressive effects on translation (Hara *et al*. 2018). Indeed, translation of hundreds of transcripts are mis-regulated in the absence of PNG activity (Kronja *et al*. 2014b). Coupled with the fact that PNG kinase activity is limited to a precise developmental time window (Hara *et al*. 2017), this widespread regulation of translation serves to coordinate multiple essential processes needed for progression through the MZT.

## Data Availability

Strains and plasmids are available upon request. Sequencing data have been deposited in GEO under accession code GSE197582. Reviewers can access the data using token **erqlmeiqljmzvyd**. Differentially expressed gene lists can be found in Supplemental Table 1 and lists of reagents can be found in the Reagents Table.

## Acknowledgements

We acknowledge Howard Lipshitz and Craig Smibert for helpful discussions and providing fly stocks. We thank Terry Orr-Weaver for providing the *twe^HB5^ cn bw/CyO* stock, Bloomington Stock Center for *Drosophila* stocks and the *Drosophila* Genomics Resource Center, supported by NIH grant 2P40OD010949 for the UASt-attB plasmid. This work was supported by the Northwestern University NUSeq Core Facility, which performed the high-throughput sequencing.

E.D.L., H.K., C.Y.L., and M.M.H. conceived the study. E.D.L., H.K., Z.A.F., S.D.K and M.M.H. performed the experiments. E.D.L., H.K., Z.A.F., S.D.K., C.Y.L. and M.M.H. analyzed the data. E.D.L. and M.M.H. wrote the manuscript.

---

**Supplemental Table 1: Lists of differentially expressed genes.** Lists of differentially expressed genes at each time point collected (late stage oocyte, NC10, NC12, NC13, early NC14 and later NC14) between the *brat* mutant and *HisRFP* embryos including the log2(foldchange,) p value, adjusted p value, MZT class, whether or not the gene is bound by ZLD, whether or not the gene is bound by BRAT, whether or not the gene was upregulated in Laver *et al*. 2015, the zygotic class, whether or not the gene was considered zygotically expressed by Strong *et al*. 2020 and whether or not the gene determined to be differentially expressed using the criteria log_2_(fold change) >1, padj <0.05. Comparisons between the *brat* hets and *HisRFP*, and clusters from Figure 5A and classes from Figure 5C are also included.

**Supplemental Figure 1:**
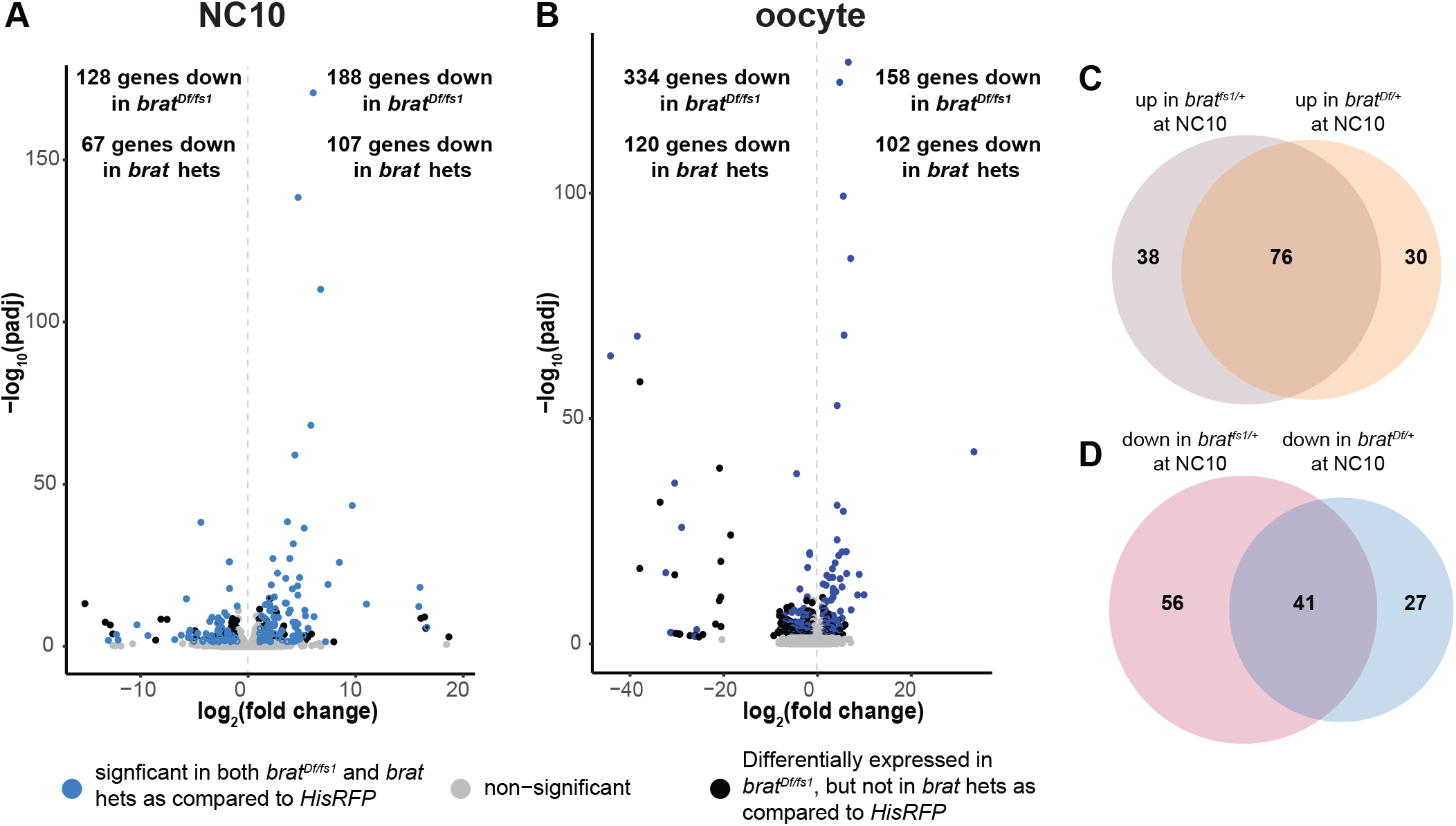
Embryos laid by mothers heterozygous for *brat*-mutant alleles (*brat* het) have differences in gene expression as compared to controls. A. Volcano plot of differentially expressed genes (log_2_fold change >1, padj <0.05) in NC10 embryos laid by *brat^Df/fs1^* mothers as compared *HisRFP* NC10 embryos. Genes that are also misexpressed in NC10 embryos laid by mothers heterozygous for a *brat*-mutant allele (*brat* het) as compared to *HisRFP* cotrols are in blue. All other significantly differentially expressed genes are in black and non-significant genes are in gray. B. Volcano plot of differentially expressed genes (log_2_fold change >1, padj <0.05) in *brat^Df/fs1^* oocytes compared *HisRFP* oocytes. Genes that are also misexpressed in *brat* het oocytes compared to *HisRFP* cotrols are in blue. All other significantly differentially expressed genes are in black and non-significant genes are in gray. C. Overlap of individual genes that show increased expression in NC10 embryos laid from either *brat^fs1/+^* or *brat^Df/+^* heterozygous mothers. D. Overlap of individual genes that show decreased expression in NC10 embryos laid from either *brat^fs1/+^* or *brat^Df/+^* heterozygous mothers.

**Supplemental Figure 2:**
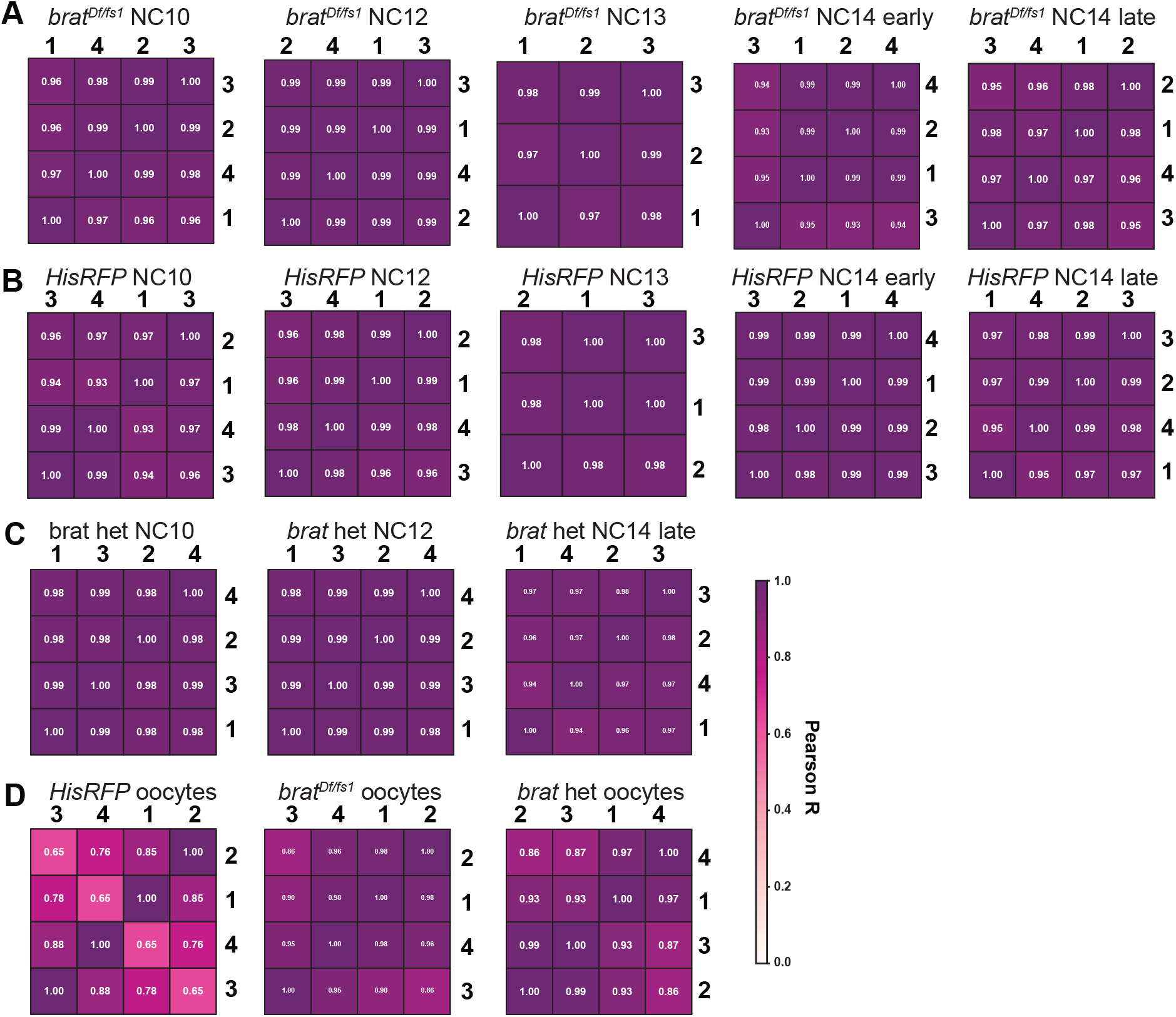
Single-embryo and single-oocyte replicates are highly correlated. A-D. Correlation plots of replicates from RNA-seq experiments in *bratD^f/fs1^* NC10-NC14 embryos (A), *HisRFP* NC10-NC14 embryos (B), *brat* heterozygous (*brat* het) NC10-NC14 embryos (C) and oocytes (D). The color of each square represents the Pearson R coefficient. Replicates numbers are listed.

